# Improving tuberculosis surveillance by detecting international transmission using publicly available whole-genome sequencing data

**DOI:** 10.1101/834150

**Authors:** Andrea Sanchini, Christine Jandrasits, Julius Tembrockhaus, Thomas Andreas Kohl, Christian Utpatel, Florian P. Maurer, Stefan Niemann, Walter Haas, Bernhard Y. Renard, Stefan Kröger

**Author notes:** Equal contribution (Andrea Sanchini, Christine Jandrasits). **Corresponding author information** Bernhard Y. Renard, Hasso Plattner Institute, Faculty for Digital Engineering, University of Potsdam, Potsdam, Germany. **Conflict of interest:** None declared.

## Abstract

**Introduction:** Improving the surveillance of tuberculosis (TB) is especially important for multidrug-resistant (MDR) and extensively drug-resistant (XDR)-TB. The large amount of publicly available whole-genome sequencing (WGS) data for TB gives us the chance to re-use data and to perform additional analysis at a large scale.

**Aim:** We assessed the usefulness of raw WGS data of global MDR/XDR-TB isolates available from public repositories to improve TB surveillance.

**Methods:** We extracted raw WGS data and the related metadata of Mycobacterium tuberculosis isolates available from the Sequence Read Archive. We compared this public dataset with WGS data and metadata of 131 MDR- and XDR-TB isolates from Germany in 2012-2013.

**Results:** We aggregated a dataset that includes 1,081 MDR and 250 XDR isolates among which we identified 133 molecular clusters. In 16 clusters, the isolates were from at least two different countries. For example, cluster2 included 56 MDR/XDR isolates from Moldova, Georgia, and Germany. By comparing the WGS data from Germany and the public dataset, we found that 11 clusters contained at least one isolate from Germany and at least one isolate from another country. We could, therefore, connect TB cases despite missing epidemiological information.

**Conclusion:** We demonstrated the added value of using WGS raw data from public repositories to contribute to TB surveillance. By comparing the German and the public dataset, we identified potential international transmission events. Thus, using this approach might support the interpretation of national surveillance results in an international context.

## Introduction

Improving the surveillance of Tuberculosis (TB) is one of the eight core activities identified by the World Health Organization (WHO) and the European Respiratory Society to achieve TB elimination, defined as less than one incident case per million [1]. Monitoring transmission is especially important for multidrug-resistant (MDR)-TB isolates – defined as being resistant to rifampicin and isoniazid – and for extensively drug-resistant (XDR)-TB isolates – defined as MDR-TB isolates with additional resistant to at least one of the fluoroquinolones and at least one of the second-line injectable drugs. In 2017, the WHO estimated that worldwide more than 450,000 people fell ill with MDR-TB and among these, more than 38,000 fell ill with XDR-TB [2].

The rapid advance in molecular typing technology – especially the availability of whole-genome sequencing (WGS) to identify and characterize pathogens – gives us the chance of integrating this information into the disease surveillance. For TB surveillance it is possible to combine the results of molecular typing of *Mycobacterium tuberculosis* complex isolates with traditional epidemiological information to infer or to exclude TB transmission [3,4]. This is of particular relevance if transmission occurs among multiple countries, where epidemiological data such as social contacts are more difficult to get and where data exchange is more difficult to organize. The European Centre for Disease Prevention and Control (ECDC) identified 44 events of international transmission (international clusters) of MDR-TB isolates collected in different European countries between 2012 and 2015 [5]. In this example, the authors inferred TB transmission using the mycobacterial interspersed repetitive units variable number of tandem repeats (MIRU-VNTR) typing method. However, this method has limitations such as low correlation with epidemiological information in outbreak settings and low discriminatory power [3,6]. In comparison, WGS analysis offers a much higher discriminatory power and allows for inferring (or excluding) TB transmission at a higher resolution [4]. In a recent systematic review, van der Werf and co-authors identified three studies that used WGS to investigate the international transmission of TB [7].

In recent years, the amount of WGS data available is increasing, especially due to the reduction of sequencing costs [8]. In addition, more and more authors deposit the raw data of their projects in open access public repositories such as the Sequence Read Archive (SRA) of the National Center for Biotechnology Information (NCBI) [9]. These raw WGS data of thousands of isolates – together with their public availability – enable the re-use and the additional analysis at a large and global scale [10]. For example, it is possible to compare genomic data among different studies/countries since the data are available in a single place. Moreover, new software/tools can be tested using the same raw WGS data [11]. However, standards in bioinformatics analysis and interpretation of these WGS data for surveillance purposes are not yet fully established [12].

We aimed to assess the usefulness of raw WGS data of global MDR/XDR-TB isolates available from public repositories to improve TB surveillance. Specifically, we wanted to identify potential international events of TB transmission and to compare the international isolates with a collection of *M. tuberculosis* isolates collected in Germany in 2012-2013.

## Methods

### Data collection: public dataset

The SRA database is a public repository provided by the NCBI (U.S. National Library of Medicine, Bethesda, USA) which stores raw sequencing data derived from high-throughput sequencing platforms [9]. We queried the repository for the pathogen “*Mycobacterium tuberculosis*” and restricted the results to “*genomic*”, “*WGS*” data from the “*Illumina*” sequencing technology using the appropriate query keywords. After excluding single-end sequenced and missing raw data, 8,716 isolates remained, which were further filtered for sequence characteristics. We excluded samples with reads shorter than 100 bp, as well as samples with a relatively low (< 20x) or high (> 500x) average coverage depth of the reference genome (see below) to obtain a more homogenous dataset. In addition, we excluded samples with less than 90% reads aligned to the reference genome to prevent having contaminated or incorrectly annotated samples in the set. Samples for which over 50% of all single-nucleotide variant calls were inconclusive were also excluded (see Supplementary Material for details). To identify duplicates (e.g. the same file uploaded more than once in different projects) within the public dataset, we compared numbers of reads and detected variants at every step of the analysis. We excluded samples that were identical in all those numbers and their corresponding epidemiological data. After all filtering steps, 7,620 isolates remained and we will refer to these isolates as the “public dataset” throughout the manuscript. In addition to the raw reads, we also collected metadata available in the SRA repository [9] (for details see Supplementary Table S1).

### Data collection: German dataset

In addition to the international public dataset, we analyzed isolates from Germany, which will be referred to as “German dataset” throughout the manuscript. The German dataset includes all *M. tuberculosis* complex isolates processed at the National Reference Center for Mycobacteria (*Forschungszentrum Borstel*, Germany) and classified as MDR-TB or XDR-TB in 2012-2013 by drug susceptibility tests (DST) according to the German TB surveillance system [13]. We extracted the epidemiological data available for the *M. tuberculosis* complex isolates using the laboratory ID of the National Reference Center for Mycobacteria. Then, we identified the respective isolate in the national surveillance system at the Robert Koch Institute (the German public health institute) and thus matched molecular with epidemiological data. We collected information on year of isolation, federal state of isolation, DST results, and patient-related information such as age, sex, citizenship, and country of birth.

### Ethical statement

Ethical approval was not required for this study since data were extracted from pseudonymized notification data.

### NGS analysis workflow

Raw reads were subjected to quality control with Trimmomatic [14] and Flash [15]. The trimmed and filtered reads were mapped to two different reference genomes: the *M. tuberculosis* H37Rv strain and a pan-genome reference built from 146 *M. tuberculosis* genomes [16,17] with bwa mem [18]. Duplicated reads were marked and reads with mapping quality less than 10 were excluded. The Genome Analysis Toolkit (GATK) [19] was used for variant detection mapped to both reference genomes and extracted all SNPs of high quality (see Supplementary Material for details).

### Drug-resistance prediction

We used Phyresse [20] and TBDreamDB [21] to identify drug-resistance mutations in our datasets (last access October 18^th^, 2018). We filtered both lists to include only single nucleotide substitutions. For TBDreamDB we mapped the provided locations within resistance genes to positions on the *M. tuberculosis* H37Rv genome where necessary. We excluded mutations not associated with drug-resistance according to the WHO [22] and to the CRyPTIC study [23] (see Supplementary Table S2 for the list of all identified mutations and, among those, all the excluded mutations). We intersected this list of mutations with the variants detected from reads mapped to the *M. tuberculosis* H37Rv genome from each sample to identify resistance-associated mutations within samples. We also identified uncovered or low-quality regions that overlap with locations of resistance mutations. For the classification of isolates into resistance classes (MDR-TB and XDR-TB), we used the definitions of the WHO [2].

### Molecular clustering

We used PANPASCO [17] to calculate relative pairwise SNP distance between all isolates classified as MDR-TB or XDR-TB in the public and German dataset. This method builds on two parts to enable distance calculation for large, diverse datasets: mapping all reads to a computational pan-genome including 146 *M. tuberculosis* genomes and distance calculation for each individual pair of samples. For this, we identified all positions with high quality for each pair of samples and calculated the SNP distance based on this set of positions (for details on the filtering workflow, PANPASCO and distance calculation see Supplementary Material). SNPs in repeat-rich genes were not used for distance calculations as studies have shown that variants found in these regions are often false positives [3,24]. The list of genes provided by Comas et al. [25] was used for filtering.

We applied single-linkage agglomerative clustering for defining transmission clusters and used a threshold of fewer than 13 SNPs, based on a previous study [26]. We chose the “upper” threshold of 13 SNPs (or >12SNPs) because we aimed to identify larger events of international transmission of TB, in contrast to the “lower” threshold of 5 SNPs, which might be more useful to identify recent transmission of TB [27,28]. Besides, we chose the “upper” threshold of 13 SNP because our isolates were spread in terms of location and time (see below) and because we are probably missing several intermediary isolates (and cases) in our collection. PANPASCO calculates distances based on data available for each pair separately. For this reason, an individual sample can potentially have small distances to samples that have a much greater distance in direct comparison, due to a higher number of compared high-quality sites. In this study, we aimed to discover clusters of closely related samples. Therefore, the implemented agglomerative clustering approach evaluates the distance from the sample that should be added to two instead of one sample of an existing cluster – we did not only compare pairs of samples but two sets of trios. The sample was added to the cluster only if the maximum distance in the trio was below twice the SNP threshold. Samples that violated this condition were iteratively removed from the clustering and were marked for potential follow-up analyses.

We used Cytoscape 3.7 to visualize the clusters [29]. We classified all clustered samples into TB lineages using lineage-specific SNPs provided in [30] and [31] (see Supplementary Table S6). We compared and validated clustering results of a subset of isolates using the pipeline MTBSeq [32] (see Supplementary Table S7).

## Results

### Final dataset

After the filtering steps, 7,620 of initially 8,716 downloaded isolates remained in the public dataset and 131 isolates from the German dataset (Figure 1). We focused our study on MDR/XDR-TB, and therefore the final dataset contained overall 1,335 isolates after filtering using resistant associated SNPs. Supplementary Table S1 shows the cluster assignment, molecular drug-resistance prediction and extracted metadata of these 1,335 isolates.

**Figure 1.**
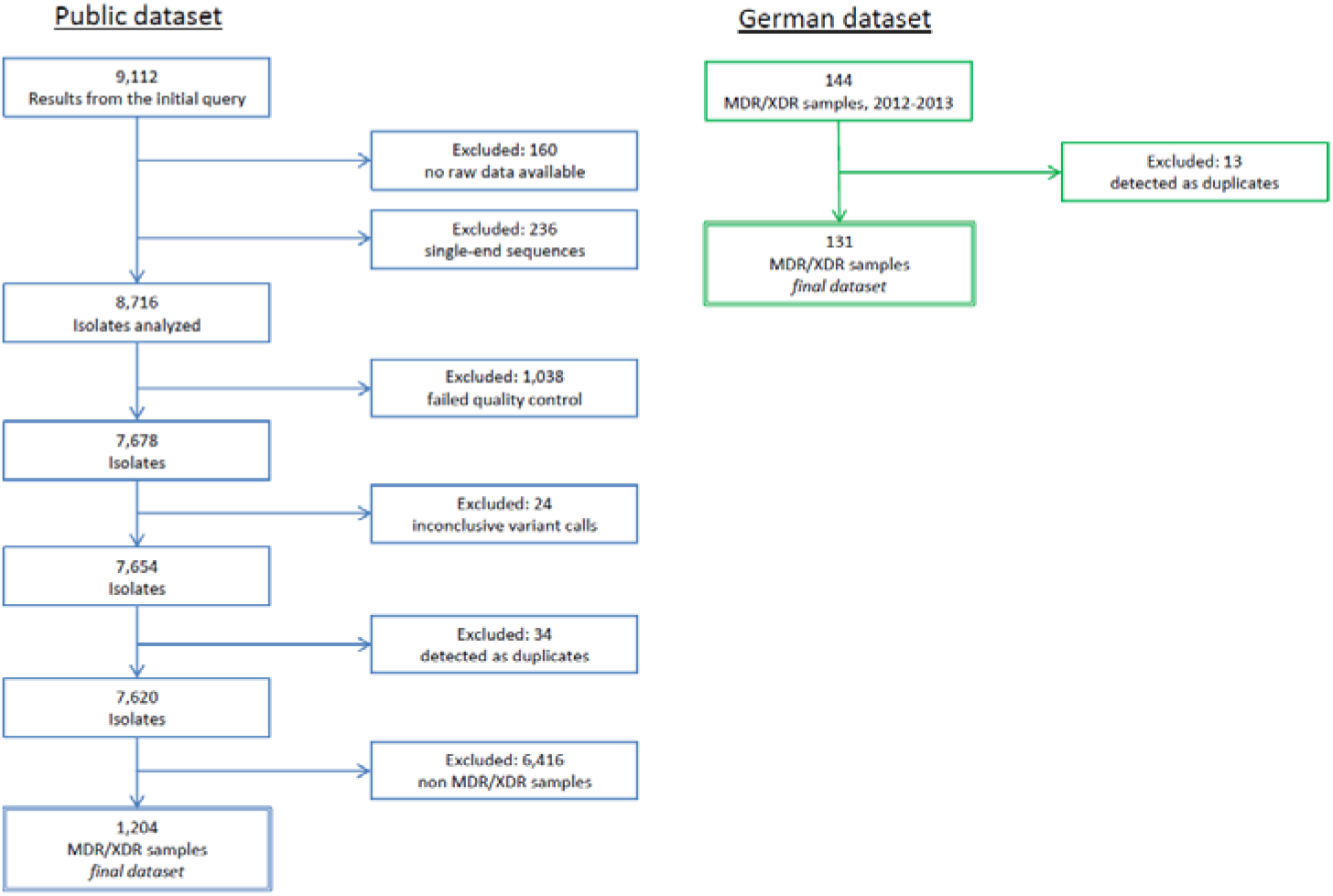
Flowchart of the inclusion and exclusion of isolates in our study from the public and the German dataset. The final dataset included 1,335 isolates: 1,204 from the public and 131 from the German dataset.

### Metadata availability and drug-resistance prediction: public dataset (N=1,204)

The majority of metadata collected from the public dataset consisted of the country of isolation (1,049/1,204, 87.13 %), the year of isolation (921/1,204, 76.49 %) and the source of the isolate (997/1,204, 82.81 %) (Table 1). For other metadata we could collect less information, for example in the case of patient age (174/1,204, 14.45 %), patient sex (171/1,204, 14.20 %), or patient HIV status (157/1,204, 13.04 %) (Supplementary Table S1). For 912 isolates, we had information on both country and year of isolation. Initially, we identified 336 isolates with missing data for the country of isolation. After examining the Bioproject information (SRA, [9]) of these 332 isolates, we could further identify the country of isolation of 177 isolates, leaving us with 155 isolates without any information regarding the country of isolation. We identified 970/1,204 MDR (80.56 %) and 234/1,204 XDR (19.44 %) isolates.

**Table 1.** Characteristics of the 1,204 multi- and extensively drug-resistant *Mycobacterium tuberculosis* isolates from the public dataset analyzed in this study.

| Characteristic |  | n |
| --- | --- | --- |
| Country of isolation | South Africa | 295 |
|  | Georgia | 160 |
|  | Moldova | 135 |
|  | Vietnam | 68 |
|  | Azerbaijan | 57 |
|  | Bangladesh | 46 |
|  | Romania | 37 |
|  | Djibouti | 31 |
|  | Ivory Coast | 29 |
|  | India | 28 |
|  | Nigeria | 27 |
|  | Thailand | 24 |
|  | Peru | 23 |
|  | China | 23 |
|  | Tanzania | 17 |
|  | Other | 49 |
|  | NA | 155 |
| Year of isolation | 2016 | 53 |
|  | 2015 | 254 |
|  | 2014 | 106 |
|  | 2013 | 147 |
|  | 2012 | 86 |
|  | 2011 | 60 |
|  | 2010 | 87 |
|  | 2009 | 65 |
|  | 2008 | 27 |
|  | 2007 | 11 |
|  | 2006 | 6 |
|  | 2005 | 14 |
|  | 2004 | 6 |
|  | 2003 | 1 |
|  | 1996 | 1 |
|  | NA | 280 |
| Source of the isolate | Sputum | 833 |
|  | Morgue | 167 |
|  | Other | 6 |
|  | NA | 198 |

NA: not available

### Metadata availability and drug-resistance prediction: German dataset (N=131)

We could retrieve demographics, epidemiological information and DST results for 129/131 (98.47 %) of the isolates from the German TB surveillance system. Table 2 and Supplementary Table S3 show the collected metadata. The 131 German isolates came from 15/16 (93.75 %) of the German federal states. The most frequent countries of birth of the patients were Russia (27/131, 20.61 %), Germany (19/131, 14.50 %) and Romania (10/131, 7.63%) (Table 2).

**Table 2.** Characteristics of the 131 multi- and extensively drug-resistant *Mycobacterium tuberculosis* isolates from Germany analyzed in this study. We found demographic information, epidemiological information and drug susceptibility test-results in the German TB surveillance system for 129/131 isolates.

| Characteristic |  | n |
| --- | --- | --- |
| Molecular drug resistance prediction | MDR | 111 |
|  | XDR | 16 |
|  | Non MDR non XDR | 4 |
| Phenotypic drug resistance prediction | MDR | 122 |
|  | XDR | 7 |
|  | NA | 2 |
| Year of isolation | 2013 | 80 |
|  | 2012 | 50 |
|  | 2014 | 1 |
| Federal state of isolation | North Rhine- Westphalia | 32 |
|  | Bavaria | 13 |
|  | Baden- Württemberg | 15 |
|  | Saxony | 10 |
|  | Lower Saxony | 10 |
|  | Berlin | 10 |
|  | Hamburg | 8 |
|  | Hesse | 8 |
|  | Schleswig- Holstein | 5 |
|  | Saxony- Anhalt | 5 |
|  | Other | 11 |
|  | NA | 4 |
| Patient age | Median | 34 (2-83) |
|  | Mean | 35.73 |
| Patient sex | Male | 79 |
|  | Female | 50 |
|  | NA | 2 |
| Patient citizenship | Germany | 30 |
|  | Russia | 25 |
|  | India | 8 |
|  | Georgia | 7 |
|  | Romania | 7 |
|  | Kazakhstan | 6 |
|  | Ukraine | 5 |
|  | other | 39 |
|  | NA | 4 |
| Patient country of birth | Russia | 27 |
|  | Germany | 19 |
|  | Romania | 10 |
|  | Ukraine | 8 |
|  | India | 8 |
|  | Kazakhstan | 8 |
|  | Georgia | 7 |
|  | Other | 41 |
|  | NA | 3 |

MDR: multidrug-resistant; XDR: extensively drug-resistant; NA: not available

We identified discrepancies in the identification of rifampicin resistance between the results of the phenotypic DST and the detection of drug-resistance mutations in 13 isolates (Supplementary Table S3). Specifically, four isolates were classified as MDR in the TB surveillance system (isolates 4556-12, 9165-12, 72-13 and 14102-13) while they were classified as non-MDR according to the molecular analysis, due to the absence of any drug-resistance mutations against rifampicin. However, in one of these four isolates (isolate 72-13), we found insufficient sequencing coverage in some of the genomic regions with known resistance mutations for rifampicin; while in another isolate (isolate 14102-13) we found an insertion of 3 nucleotides near a region with known resistance mutations for rifampicin. In addition, nine isolates were classified as MDR in the TB surveillance system (isolates 11355-13, 2955-12, 3007-13, 4245-13, 5096-13, 5190-13, 7712-13, 8291-13 and 8565-12), while they were classified as XDR according to the analysis of the drug-resistance mutations. The reason for such discrepancy was that a drug-resistance mutation against amikacin, kanamycin or capreomycin was identified in these ten isolates, but no DST results were available for these antibiotics.

### Molecular clustering and comparison between the public and the German dataset

Among all the isolates of our study, we identified 133 molecular clusters – with at least 2 isolates – and 591 singletons. The 133 clusters included 744 isolates (Supplementary Table S4). Supplementary Table S5 shows a summary of distances between all isolates for all molecular clusters. In 16 clusters, the isolates were from at least two different countries of isolation, suggesting larger events of international transmission of TB (Supplementary Table S4). For example, cluster2 included 56 MDR/XDR isolates from three countries – Moldova, Georgia and Germany. A total of 51/56 isolates in this cluster were part of a previous study (Bioproject PRJNA318002, [33], Supplementary Table S1). In Figure 2 we show the country of isolation and the year of isolation of the isolates belonging to cluster2.

**Figure 2.**
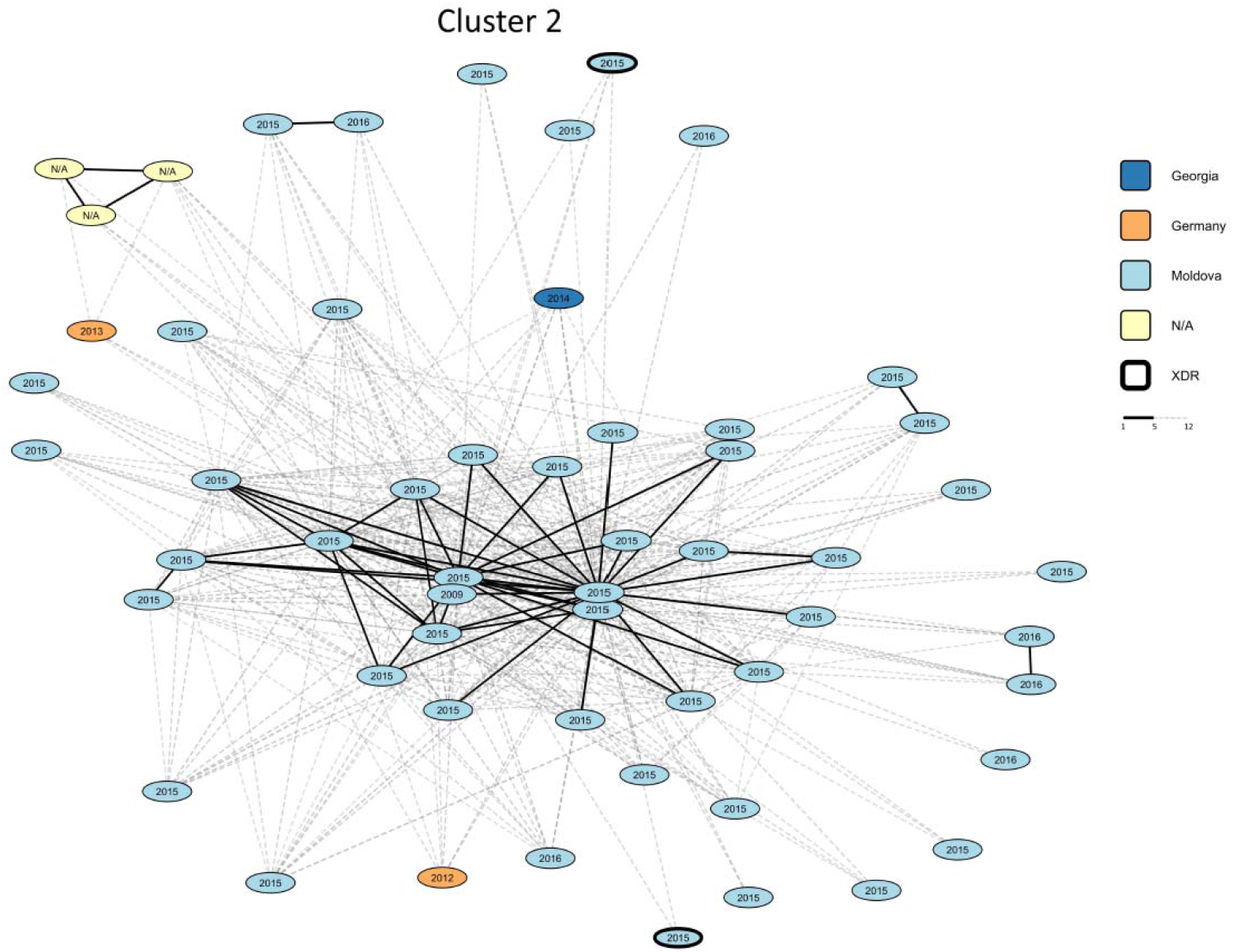
Visualization of the transmission cluster2 (N=56) identified among the 1,335 *Mycobacterium* tuberculosis isolates analyzed in our study. The country of isolation, multi- and extensive drug-resistance classification and year of isolation are represented in the clusters. SNP distances were calculated for each pair of isolates individually. Links with less than 6 SNPs are marked black, those with less than 13 SNPs are marked in grey. Connections with 13 SNPs or more than are not shown in the network.

Cluster1 is the largest cluster (n=79) identified in our study. According to the metadata (such as host subject, isolate name, year of isolation, patient age, and patient sex, see Supplementary Table S1), the isolates were 79 autopsy samples from different anatomic sites (such as lung or liver) of the same patient, marked as “P21”. Similarly, cluster3, cluster14, cluster16, cluster18 and cluster28 contained multiple isolates from single patients from South Africa, which were part of a study including 2,693 autopsy samples of 44 subjects [34]. In line with previous findings [34], our analysis showed very low variability within these clusters, highlighted by the low maximum cluster distances in each of these “single-patient” clusters (Supplementary Table S5). In addition, the analysis of the respective metadata revealed that cluster26, cluster32 and cluster33 included multiple isolates from single patients. These isolates were part of a study investigating the evolution of drug-resistant TB in patients during long-term treatment [35].

When we compared the German dataset with the public dataset, we observed that in 11 clusters there was at least one isolate from Germany and at least one isolate from another country. Table 3 shows the relation between the German isolates and the international isolates from the public dataset. The epidemiological information collected from the German isolates correlates well with molecular clusters in 7/11 cases. For example, in cluster9 there were 16 isolates from Georgia and two isolates from Germany; the country of birth recorded for one of these two isolates from Germany was Georgia. Moreover, cluster24, cluster35, and cluster103 included isolates from Georgia and Germany, and the country of birth recorded for the isolates from Germany was Georgia. Three further examples of agreement between molecular and epidemiological data were: the cluster13, which included isolates from Germany and Kazakhstan, the cluster53, which included isolates from Germany and from Romania and the cluster58, which included isolates from Germany and India (Table 3). By comparing the molecular data of the German and of the public dataset, we could connect previously epidemiologically unlinked cases. For example, in the cluster2 (Figure 2) two isolates from Germany (in orange) were connected through several isolates from Georgia and Moldova (in dark and light blue), and the distance between the two German isolates was >13 SNPs. Similarly, in the cluster53 two isolates from Romania were connected through a German isolate, and the distance between the two isolates from Romania was > 13 SNPs (data not shown).

**Table 3.**
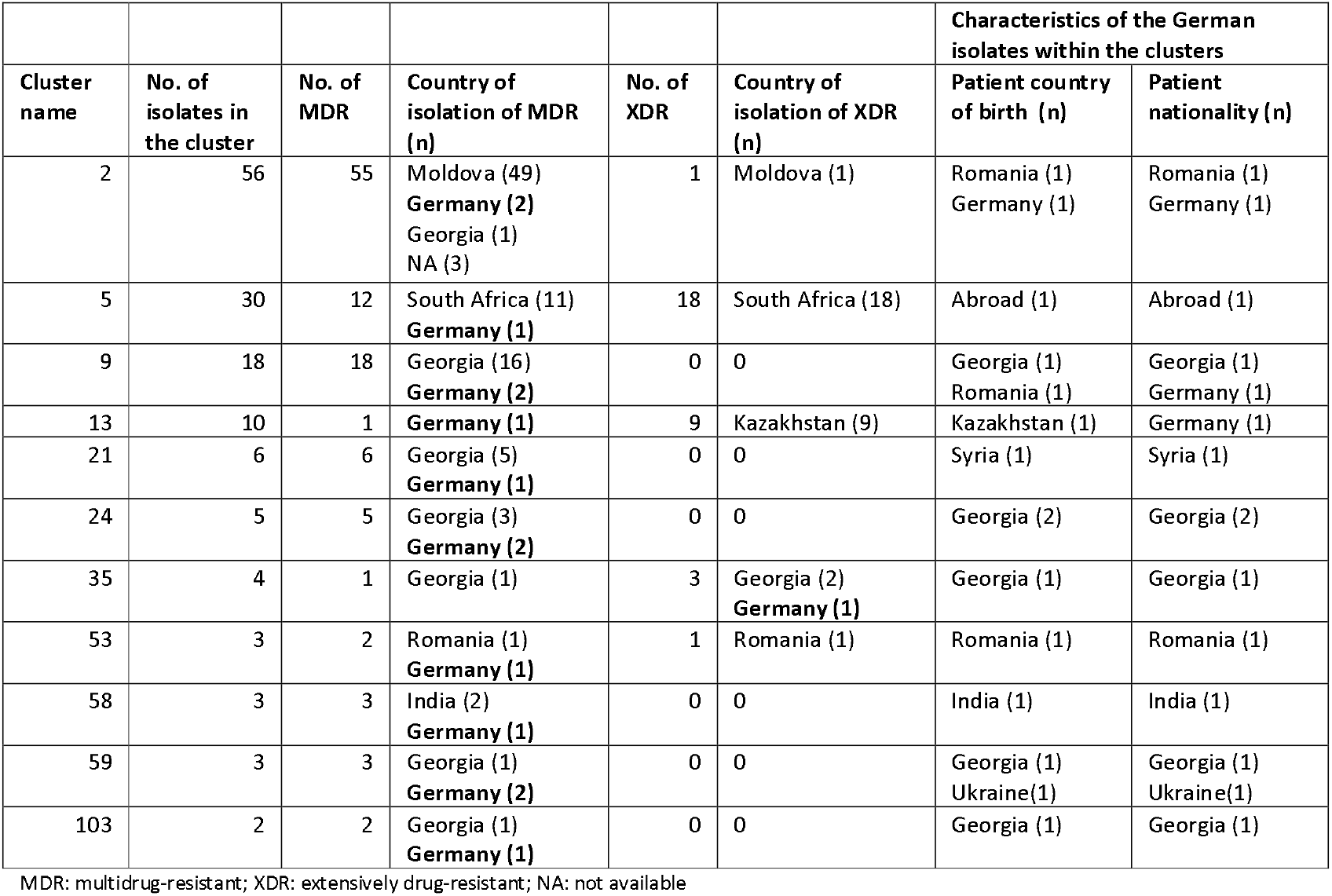
Characteristics of the 11 molecular clusters identified in this study which contain at least one isolate from Germany and at least one isolate from another country. In bold the isolates from Germany. Within each cluster, information about the country of birth, the nationality and the federal state of isolation of the German isolates is provided.

## Data availability

The raw whole genome sequencing data used in this study are available in the NCBI SRA repository. The accession numbers for all samples of the public dataset are available in the Supplementary Table 1. The German dataset is available as Bioproject PRJEB35201. Software for creating a pan-genome sequence (seq-seq-pan) is accessible at https://gitlab.com/rki_bioinformatics/seq-seq-pan and scripts for the NGS workflow and the SNP-distance method (PANPASCO) are available at https://gitlab.com/rki_bioinformatics/panpasco. The code for the clustering method is available at https://gitlab.com/rki_bioinformatics/snp_distance_clustering.

## Discussion

In this study, we assessed the usefulness of raw WGS data of MDR/-XDR-TB isolates available from public sequence repositories to improve TB surveillance. We identified several molecular clusters including isolates from multiple countries, suggesting larger events of international transmission of TB. We expected to find international TB-transmission events, also considering previous studies reporting cross-border molecular clusters [5,7]. Looking at the collected metadata, we identified several clusters with multiple isolates from the same patient or multiple autopsy samples collected from the same patient [34,35]. This shows the importance of providing complete metadata together with the publicly available molecular data. Based on the metadata, we could distinguish between clusters of isolates taken from different patients – the “real” transmission clusters – and clusters of isolates taken from a single patient. The real transmission clusters are crucial for the routine TB surveillance, while the clusters of isolates taken from the same patient are useful to study the intra-host variability of isolates.

We observed agreement between molecular and epidemiological data by comparing the public and the German datasets. This is clear for example in the clusters containing isolates from both the German dataset and the public dataset originating from Georgia. It is therefore likely that migrants from Georgia acquired the TB infections in their country – or during visits there – and were diagnosed later when they moved or returned to Germany, as already described [36]. This shows that we could identify events of potential international transmission (between Germany and Georgia), that we could have missed by looking only at the German molecular clusters. Our analysis can have implications for surveillance and public health, for example by linking TB patients from different countries during contact tracing procedures. In the best scenario, where we could compare the German and the public dataset in (almost) real-time, we would detect international transmission of TB earlier and inform the public health authorities timely.

We observed discrepancies in the identification of rifampicin resistance between the results of the phenotypic DST and the detection of drug-resistance mutations. Specifically, four isolates were phenotypically resistant to rifampicin but they did not contain any known drug-resistance mutation against rifampicin or the genetic regions containing the known mutation had lower sequencing quality. This means that in our study the known drug-resistance mutations (there might be always new mutations conferring resistance that arise) correctly predicted the resistance to rifampicin in 125/129 of the isolates, resulting in a sensitivity of 96.90 %. This sensitivity is in accordance with a study by the CRyPTIC Consortium, where the authors reported a sensitivity of 97.50 % [23]. Misclassification occurred in four isolates, which were MDR by phenotype, but non-MDR by genotype. This might have had consequences for patient therapy if we would have replaced the phenotypic DST with the molecular detection of drug-resistance mutations. Therefore, we suggest being careful in the transition from phenotypic to genotypic drug-resistance determination as suggested by the CRyPTIC Consortium [23]. Specifically, laboratories and national reference laboratories should still perform the phenotypic DST, for example on a representative set of isolates or isolates with low sequencing quality and coverage.

Our study has three major limitations: first, the raw WGS data uploaded in the SRA repository [9] were either from single studies or from outbreaks, and therefore they were not representative of the TB situation in the different countries. Besides, we are probably missing several intermediary isolates (and cases) in our collection. These examples of sampling biases are, however, well-known biases in molecular epidemiology studies [37]. Second, the metadata collected were incomplete, especially regarding patient information. Both limitations can be overcome by genotyping all TB isolates, by including the genotyping results in the TB surveillance systems and by making genotyping data publicly available. Third, we relied on single-linkage clustering, as this is currently a widely used approach for transmission cluster detection [38]. However, with many missing samples (which is probably the case when using a public repository), single-linkage can become unreliable and results strongly dependent on the cut-off and sample coverage for the specific cluster. For our study and its exploratory purpose, we preferred to use a widely accepted cluster detection approach combined with a higher cut-off. Therefore, we preferred to err on the side of caution (by identifying a false positive connection) rather than miss a potential transmission. Thus, in future studies, researchers should carefully evaluate the clustering methods for transmission cluster detection with missing data.

In conclusion, our study has one major implication: we demonstrated that by considering the international context (the public dataset), while analysing the national molecular data (the German dataset), we could identify previously unknown transmissions between patients. Thus, we could detect larger and international events of TB transmission. To improve the WGS-based TB surveillance we, therefore, suggest to regularly compare the national molecular clusters with the international molecular clusters available in the public sequence repositories. Lastly, supranational institutions such as the WHO, the ECDC or international TB networks could perform such analysis at a global scale, improving the global surveillance of TB.

## Acknowledgements

We would like to thank Birgit Voß for her 318 help in matching epidemiological and molecular data. We want to thank the National Reference Center for Mycobacteria in Borstel, Germany for acquiring the sequencing data for the German dataset. We thank Lena Fiebig and Marta Andrés for their initial input in the study.

## Authors’ contributions

Andrea Sanchini: participated in the study design, participated in the data collection, analyzed the data, interpreted the results and wrote the manuscript.

Christine Jandrasits: designed the study, collected the data, analyzed the data, interpreted the results and wrote the manuscript.

Julius Tembrockhaus: collected the data, analyzed the data and revised the manuscript.

Thomas Andreas Kohl: participated in data analysis, and interpretation of the results, and revised the manuscript.

Christian Utpatel: participated in data analysis, interpretation of the results, and revised the manuscript.

Florian P. Maurer: participated in data analysis, and revised the manuscript.

Stefan Niemann: participated in study design and revised the manuscript.

Walter Haas: designed the study, participated in the interpretation of the results and revised the manuscript.

Bernhard Y. Renard: designed the study, participated in the interpretation 340 of the results, coordinated the project and revised the manuscript.

Stefan Kröger: designed the study, participated in the data analysis, participated in the interpretation of the results, coordinated the project and revised the manuscript.

