## Supplementary Material for "Improving tuberculosis surveillance by detecting international transmission using publicly available whole-genome sequencing data"

**Description of the sequencing analysis workflow and filtering of mixed base calls used in our study.**

**Next Generation Sequencing Analysis Workflow**

Our NGS workflow includes quality control, read mapping, variant discovery and detection of low-quality regions for paired-end next generation sequencing samples. First, adapter sequences were removed from the ends of the reads with Trimmomatic [1], then they were merged with Flash [2] in case they overlap with at least 5 base pairs. This resulted in two sets of reads: non-overlapping paired-end reads and merged, longer single reads. We used Trimmomatic [1] to trim bases of low base quality in both sets. Reads with a length of less than 35 were removed from the dataset and remaining mates of excluded reads added to the set of single reads.

We mapped all reads to two reference genomes with bwa mem [3] and the sets of mapped paired-end and single reads were joined for the following analysis with samtools [4]. We used the *Mycobacterium* *tuberculosis* H37Rv strain (Accession number NC_000962.3) for resistance mutation detection and the linear pan-genome consensus sequence built from 146 *M. tuberculosis* genomes [5, 6] for SNP distance calculation.

In preparation for variant detection, duplicated reads were marked and read groups added with picard tools [7]. Reads with a mapping quality of less than 10 were excluded from the set of mapped reads. For variant detection we used the Genome Analysis Toolkit (GATK [8]) for both genomes. As described in the workflow for PANPASCO [6], we first detected confidence scores for all sites, including reference sites with the HaplotypeCaller tool. After that we extracted genotypes for each site using the GenotypeGVCFs tool. We called variants with a diploid model to be able to filter mixed base calls by allele frequency. We analyzed all sites and separated them in variant sites, positions with uncalled genotypes and high-quality reference sites. Variant sites were then split into single nucleotide polymorphisms (SNPs), small deletions and insertions and structural variants with the SelectVariants tool from GATK. We identified SNPs with an allele frequency of at least 75% and where 5 or more reads were used to call the SNP. We also used bedtools [9] to extract regions with less than 5 reads coverage.

By using these filters we separated the data into high-quality SNPs and five sets of low-quality regions with the following criteria:

- positions with less than 5 mapped reads
- positions at deletions
- positions with uncalled genotypes
- SNPs called from less than 5 reads
- SNPs with less than 75% allele frequency

**Filter samples for inconclusive variant calls**

For mixed base call detection, we extracted all single nucleotide base calls in high quality regions called from at least 5 reads. We determined the number of SNPs with an allele frequency between 25% and 75% (mixed base calls). Samples where 50% or more high quality SNPs were mixed base calls were excluded from our analysed set.
